## Supplemental Information for "Statistical learning beyond words in human neonates"

**Table S1. Voice stimuli.**

|  | **Voice** | **Gender** | **Pitch (Hz)** |
| --- | --- | --- | --- |
| **Ma** | fr3 | Male | 75 |
| **Mb** | fr1 | Male | 108 |
| **Mc** | fr7 | Male | 140 |
| **Fa** | fr2 | Female | 133 |
| **Fb** | it4 | Female | 190 |
| **Fc** | fr4 | Female | 247 |

**Table S2. Stimuli.**

| **Experiment 1** | | | | **Experiment 2** | | | |
| --- | --- | --- | --- | --- | --- | --- | --- |
| **List A** | | **List B** | | **List A** | | **List B** | |
| **Word** | **Part-word** | **Word** | **Part-word** | **Word** | **Part-word** | **Word** | **Part-word** |
| kida | dape | dape | kida | FbMb | MbFc | MbFc | FbMb |
| petu | tubo | tubo | petu | FcFa | FaMa | FaMa | FcFa |
| bogɛ | gɛki | gɛki | bogɛ | MaMc | McFb | McFb | MaMc |

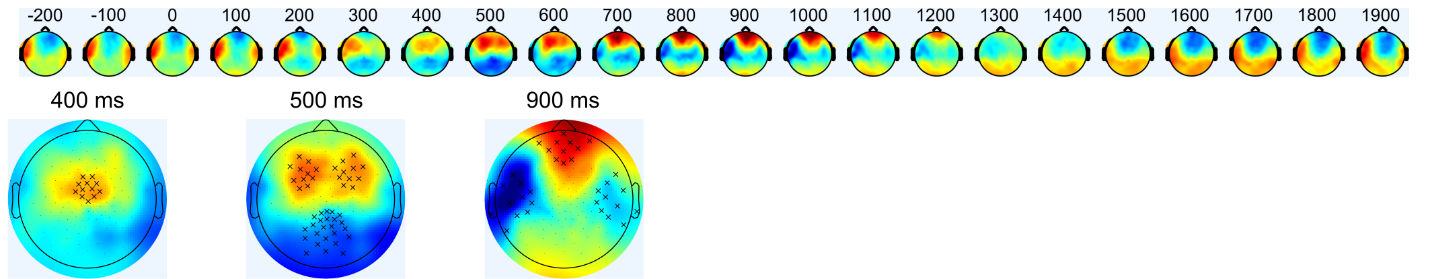

**Figure S1. Topographies for the grand average ERP**

ERP across all participants for both Experiments. The three main topographies observed during the response are plotted on the bottom. The markers show the electrodes belonging to the 7 defined ROIs: for the first topography, central electrodes; for the second topography, frontal left, frontal right and occipital electrodes; and for the third topography, temporal left, temporal right and pre-frontal electrodes. Color scale limits [-0.07, 0.07] a.u.

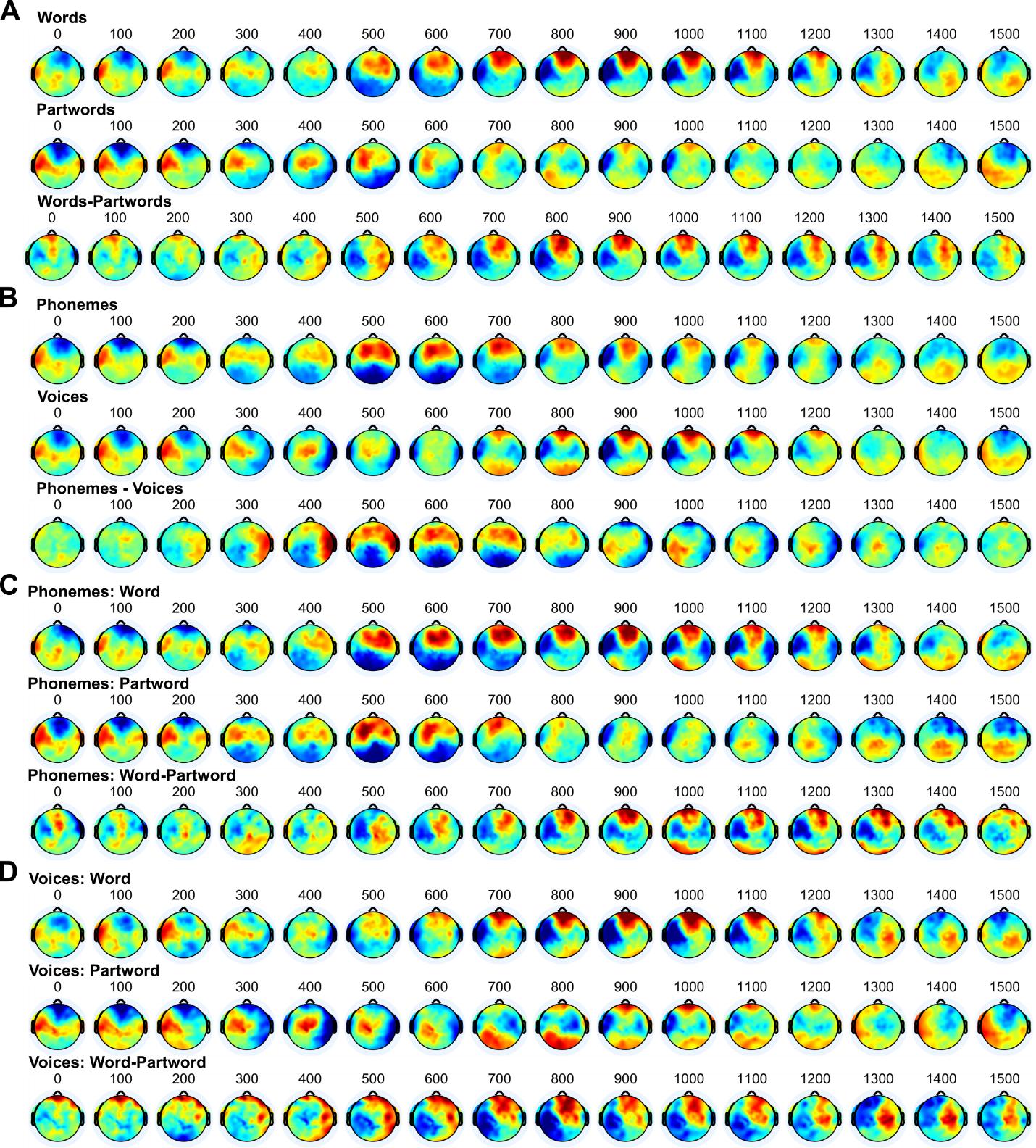

**Figure S2. Topographies for the ERPs to test words during recognition**

Color scale limits [-0.07, 0.07] a.u.

(A) Topographies for the Test-word effect

(B) Topographies for the Famliarisation effect

(C) Topographies for the Test-word effect during Experiment 1 (structured over Phonemes)

(D) Topographies for the Test-word effect during Experiment 2 (structured over Voices)

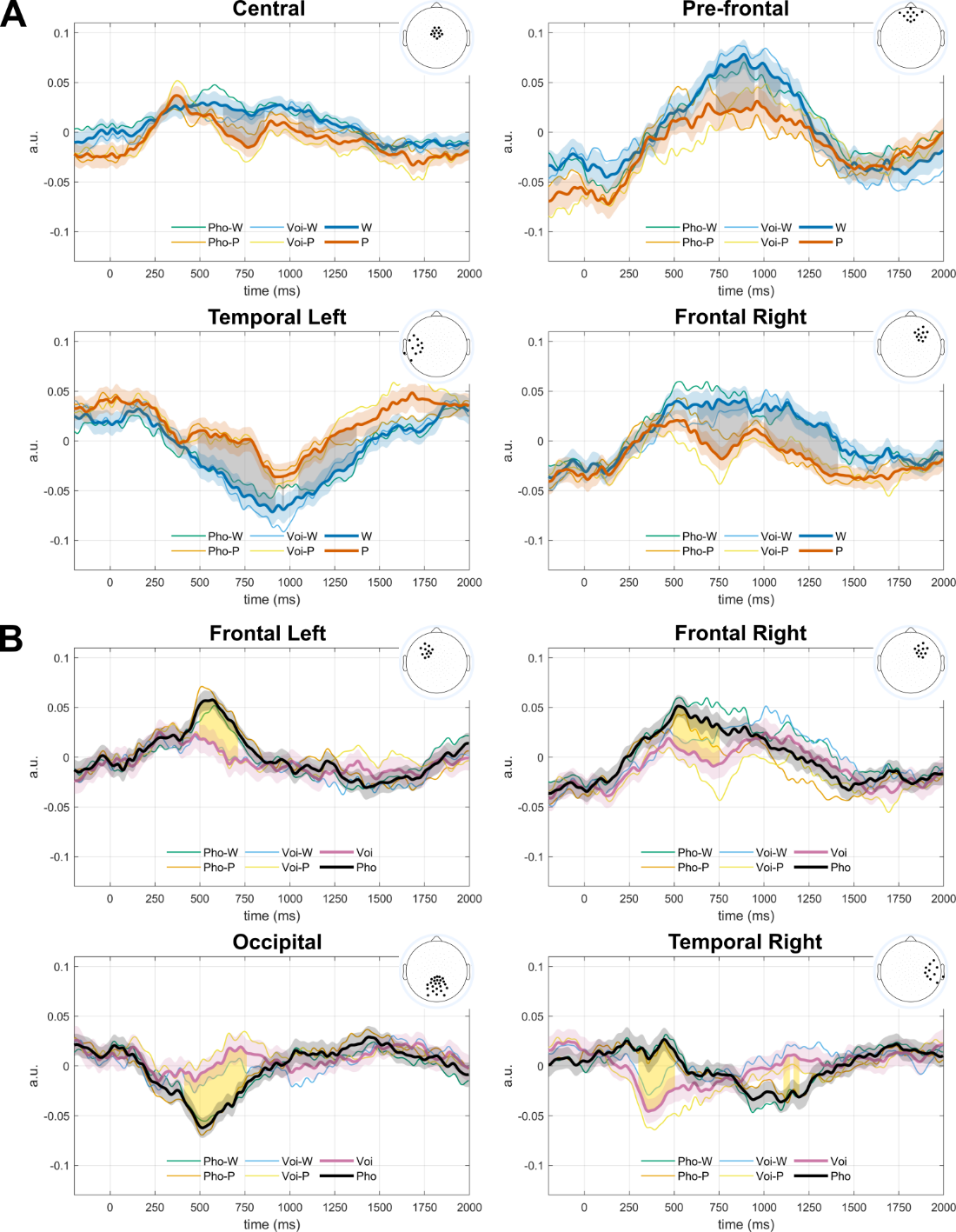

**Figure S3. Result for the ROI analysis of the ERPs to test words during recognition.**

ROIs showing significant differences for the ERPs during the recognition phase in A Words vs Partwords comparison and B Voice vs Phoneme Comparison: the thick lines show the grand averages for the two conditions showing the main effect. Shaded areas around the lines correspond to the standard error across participants. Thin lines show the ERPs separated by duplet type and Familiarization type. The shaded areas between the thick lines indicate time windows where significant differences were found after correcting by multiple comparisons (p<0.05, FDR corrected by the number of ROIs and times points). The topographies represent the electrodes belonging to the ROI. We run an ANOVA for the average activity in each ROI and significant time window, including test duplet and familiarisation as factors. We did not observe significant interactions in any case. Voi=Voice; Pho=Phoneme; W=Words, P=Part-Words.

**Post-hoc test: investigating female and male voice perception effects**

Given that gender may be an important factor in infants' speech perception (newborns, for instance, prefer female voices at birth), we conducted tests to assess whether this dimension could have influenced the results observed in Experiment 2.

*Computation of TPs*

We first quantified the transitional probabilities matrices during the structured stream of Experiment 2, considering that there were only two types of voices: Female and Male.

For List A, all transition probabilities were equal to 0.5 (P(M|F), P(F|M), P(M|M), P(F|F)), resulting in flat TPs throughout the stream (Figure S4, top). Therefore, we would not expect neural entrainment at the word rate (2 Hz) nor anticipate ERP differences between the presented duplets in the test phase.

For List B, P(M|F)=P(F|M)=0.66 while P(M|M)=P(F|F)=0.33, without a regular pattern of TP drops throughout the stream (Figure S4, bottom). Although this pattern is unlikely to induce strong neural entrainment at 2 Hz, some degree of entrainment might have occasionally occurred due to some drops occurring at a 2 Hz frequency. Regarding the test phase, all three Words and only one Part-word presented alternating patterns (TP=0.6). Therefore, we cannot rule out that gender alternation might have affected the difference in ERPs between Words and Partwords in List B.

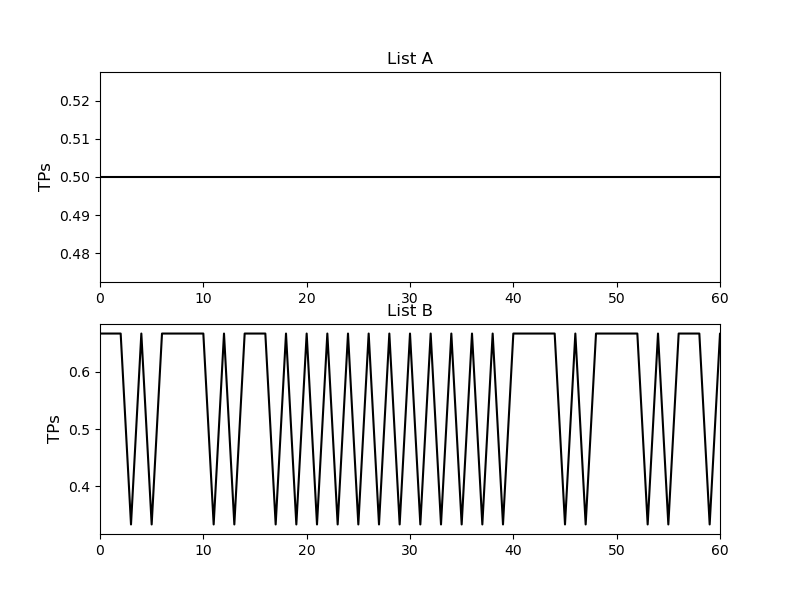

**Figure S4.** Transition probabilities (TPs) across the structured stream in Experiment 2, considering voices processed by gender (Female or Male). Top: List A. Bottom: List B.

While it seems unlikely that gender alternation alone explains the entire pattern of results, as the effect was inconsistent and appeared in only one of the lists, we separately analysed the entrainment and ERP effects in each list to rule out this possibility.

*Neural entrainment effect*

We computed the average entrainment over the electrodes, which showed significant entrainment at 2 Hz (word rate). A comparison of the 2 Hz entrainment between participants who completed List A and List B showed no significant differences (t(30) = -0.27, p = 0.79). A test against zero for each list indicated significant entrainment in both cases (List A: t(17) = 4.44, p = 0.00036; List B: t(13) = 3.16, p = 0.0075). See Figure S5.

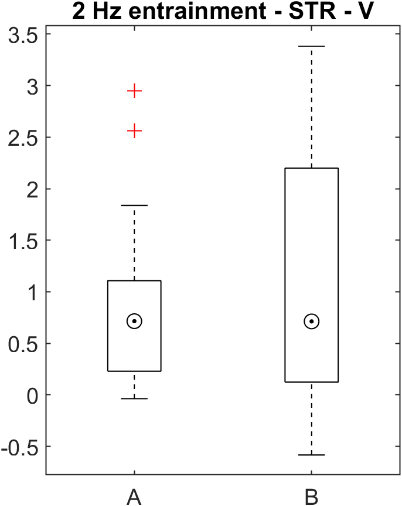

**Figure S5.** Neural entrainment at word rate (2Hz) during the structured stream of Experiment 2 for Lists A and B.

*ERP effect*

We computed the mean activation within the time windows and electrodes of interest and compared the effects of word type and list using a two-way ANOVA. For the difference between Words and Part-words over the positive cluster, we observed a main effect of word type (F(1,31) = 5.902, p = 0.021), with no effects of list or interactions (ps > 0.1). Over the negative cluster, we again observed a main effect of word type (F(1,31) = 10.916, p = 0.0016), with no effects of list or interactions (ps > 0.1). See Figure S6.

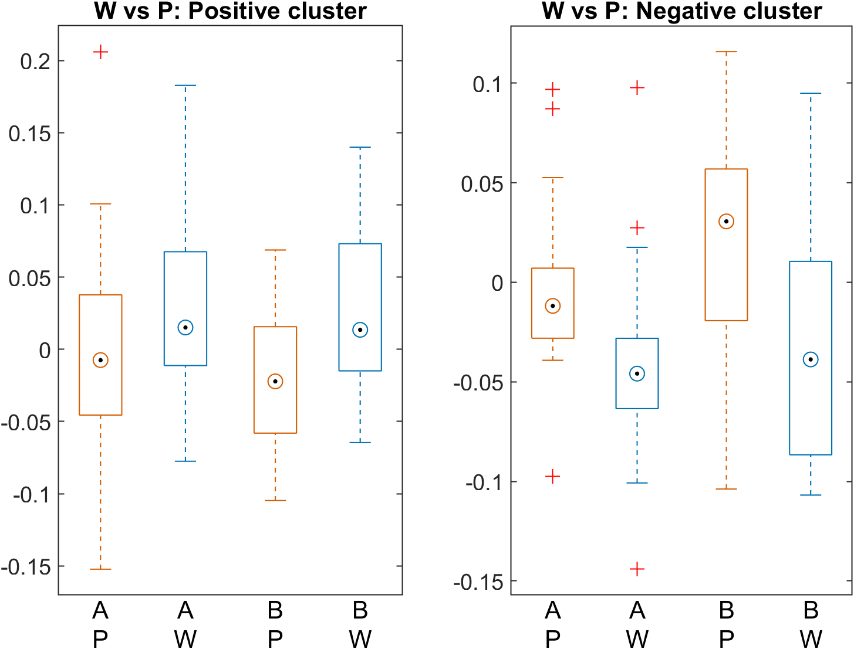

**Figure S6:** Difference in ERP voltage (Words – Part-words) for the two lists (A and B); W=Words; P=Part-Words.
